# Revisiting Logistic Regression for High-Dimensional Gene Expression Data

**DOI:** 10.64898/2026.07.20.739668

**Authors:** Rossana O. Souza, Wellington Francisco Rodrigues, Bráulio R. G. M. Couto, Marcos A. dos Santos

## Abstract

Logistic regression remains a widely used classification method due to its interpretability and computational efficiency, but its direct application to high-dimensional biomedical data is limited when the number of features greatly exceeds the number of samples. In this paper, we propose a reformulated logistic regression framework designed for feature selection and classification in complex high-dimensional settings. The method is evaluated on three biomedical datasets, including scenarios with tens of thousands of attributes and substantially fewer samples. Across these datasets, the proposed approach achieved clear separation between control and disease groups while selecting a compact set of features. Several selected features were consistent with previously reported disease-associated markers, supporting the biological plausibility of the model, while additional selected features suggest potential novel candidates for further investigation. These results indicate that the proposed framework may provide an interpretable and computationally efficient alternative for feature selection in high-dimensional computational biology applications.

## 1 Introduction

High-throughput technologies, such as microarrays and RNA sequencing, have substantially increased the dimensionality of biomedical datasets (Bir-Jmel et al., 2024; Mongardi et al., 2024). In gene expression studies, it is common to measure thousands of molecular attributes from a limited number of biological samples (Mongardi et al., 2024). This scenario, commonly referred to as the *p* ≫ *n* setting, in which the number of attributes far exceeds the number of samples, poses major challenges for statistical and machine learning methods (Mongardi et al., 2024; Borah et al., 2024). In this context, classification models are more susceptible to overfitting, unstable coefficient estimation, and poor generalization (Mongardi et al., 2024; Bir-Jmel et al., 2024).

These challenges are especially relevant in biomedical applications such as sepsis, where gene expression profiles may contain useful diagnostic information but are typically available for relatively small cohorts (Rashid et al., 2024; Scicluna et al., 2025). Beyond predictive performance, biomedical studies often require interpretable models capable of identifying candidate genes or molecular signatures associated with the condition under investigation (Mongardi et al., 2024; Scicluna et al., 2025). Therefore, methods applied to this type of data should not only distinguish between clinical groups, but also provide a meaningful and compact set of selected attributes (Mongardi et al., 2024; Lukaszuk and Krawczuk, 2024).

Logistic regression is a widely used statistical model for binary classification and remains attractive in biomedical research due to its interpretability, probabilistic formulation, and computational simplicity (Musib et al., 2024; Mahmood et al., 2025). However, the direct application of classical logistic regression to high-dimensional gene expression data is limited (Bir-Jmel et al., 2024; Mongardi et al., 2024). When the number of attributes is much larger than the number of samples, the estimation problem may become ill-conditioned, and the fitted model may suffer from instability or complete separation (Mongardi et al., 2024; Bir-Jmel et al., 2024). Regularized variants, such as LASSO and elastic net logistic regression, are commonly used to address these limitations by introducing sparsity or shrinkage in the estimated coefficients (Mongardi et al., 2024; Musib et al., 2024; Mahmood et al., 2025). Nevertheless, these approaches may select larger sets of correlated attributes or require iterative optimization procedures, which can affect interpretability and computational cost (Mongardi et al., 2024; Bir-Jmel et al., 2024).

In this work, we propose a scalable modified logistic regression framework for feature selection in high-dimensional gene expression data. The proposed formulation is based on a log-odds encoding of the binary class labels, followed by a linear formulation for estimating the coefficient vector. To improve computational feasibility, the problem is expressed through an equivalent augmented linear system that avoids the explicit inversion of dense matrices and allows the use of sparse linear algebra techniques. After estimating the coefficient vector, genes are ranked according to the absolute magnitude of their coefficients, |*α*_*i*_|, and the most relevant attributes are retained for classification and biological interpretation.

The proposed method is evaluated on three publicly available sepsis gene expression datasets: GSE12624, GSE13205, and GSE69063. Its performance is compared with LASSO logistic regression and elastic net logistic regression. The evaluation considers not only predictive metrics, such as AUC, accuracy, sensitivity, specificity, and F1-score, but also the stability and compactness of the selected gene signatures and the computational time required by each method (Lukaszuk and Krawczuk, 2024; Mongardi et al., 2024).

The main contribution of this work is not to show that the proposed method always outperforms regularized logistic regression in predictive accuracy. Instead, we investigate whether a reformulated logistic-regression-based model can provide a competitive and interpretable alternative for feature selection in high-dimensional biomedical datasets (Mongardi et al., 2024; Bir-Jmel et al., 2024). The results show that the proposed method achieves high classification performance while selecting compact gene sets across the evaluated datasets. This behavior suggests that the method may be useful when interpretability, parsimony, and computational simplicity are important considerations in gene expression analysis (Mongardi et al., 2024; Lukaszuk and Krawczuk, 2024).

The remainder of this paper is organized as follows. Section 2 reviews related work on logistic regression, feature selection, and high-dimensional biomedical classification. Section 3 describes the datasets used in this study. Section 4 discusses the limitations of classical logistic regression in high-dimensional datasets. Section 5 presents the proposed modified logistic regression framework. Section 6 reports the experimental results and comparative analysis. Finally, Section 7 presents the conclusions and future directions.

## 2 Related Work

Logistic regression has long been a fundamental tool in statistical learning and has been extensively applied to classification problems in bioinformatics (Hosmer et al., 2013; Hastie et al., 2009). Its interpretability and probabilistic framework make it particularly suitable for biomedical applications, including disease diagnosis, risk prediction, and biomarker discovery (Dreiseitl and Ohno-Machado, 2002). However, the increasing dimensionality of modern biological datasets has exposed important limitations of the classical formulation.

One of the main challenges arises in high-dimensional settings, where the number of features greatly exceeds the number of samples. In such scenarios, traditional logistic regression becomes ill-posed, often leading to overfitting and unstable coefficient estimation (Hastie et al., 2009).

In the context of gene expression analysis, several studies have explored the use of regularized logistic regression for identifying relevant biomarkers (Jiang et al., 2020; Wang et al., 2022). These methods aim to balance predictive performance with interpretability, a key requirement in clinical applications. Despite their effectiveness, these models may still struggle with highly correlated features and complex nonlinear relationships commonly found in biological data (Hastie et al., 2009).

Dimensionality reduction techniques have also been extensively used to address the challenges of high-dimensional data. Among them, singular value decomposition (SVD) and principal component analysis (PCA) are widely employed to extract latent structures and reduce noise (Hastie et al., 2009). In bioinformatics, these methods have been used both for visualization and as preprocessing steps for classification tasks (Wu and Zhang, 2020). However, relying solely on dimensionality reduction may lead to loss of important discriminative information.

More recently, hybrid approaches combining dimensionality reduction, feature selection, and classification have gained attention (Wang et al., 2022). These methods aim to leverage the strengths of each component to improve overall performance. Additionally, local and instance-based learning strategies, such as k-nearest neighbors and adaptive models, have been proposed to better capture data heterogeneity, especially in imbalanced datasets (Hastie et al., 2009).

## 3 Datasets

The GSE12624 dataset consists of gene expression profiles from polytrauma patients, comparing individuals who developed sepsis with non-septic controls. In this study, we considered 70 samples, including 36 non-sepsis samples and 34 sepsis samples. The dataset was used to evaluate the ability of the proposed method to identify a compact set of discriminative genes in a high-dimensional setting, where the number of measured attributes is much larger than the number of samples.

The GSE13205 dataset contains gene expression profiles obtained from skeletal muscle biopsy specimens of intensive care unit patients with sepsis-induced multiple organ failure and healthy controls. In this study, we considered 21 samples, including 8 control samples and 13 sepsis samples. Due to its small sample size, this dataset represents a particularly challenging scenario for classification and feature selection, making it useful for assessing the stability of the selected gene signatures under limited-sample conditions.

The GSE69063 dataset includes gene expression profiles obtained from peripheral blood samples collected from patients with sepsis and healthy controls. In this study, we considered 90 samples, including 33 control samples and 57 sepsis samples. This dataset provides a larger cohort compared with the other two datasets and was used to evaluate the performance, stability, and computational behavior of the proposed method in a high-dimensional sepsis classification task.

## 4 Logistic Regression in High-Dimensional Settings

Logistic regression is a widely used statistical model for binary classification (Hosmer et al., 2013). It maps a linear combination of the input attributes to the interval [0, 1] using the logistic function, allowing the model output to be interpreted as the probability of belonging to a given class.

One of its main advantages is interpretability, since the estimated coefficients indicate the contribution of each attribute to the predicted outcome. This is particularly relevant in biomedical applications, where identifying factors associated with a disease may be as important as classification performance (Vellido, 2020).

However, traditional logistic regression may become unstable in high-dimensional scenarios, especially when the number of attributes is much larger than the number of samples, commonly denoted as *p* ≫ *n*. In such cases, coefficient estimation is more susceptible to overfitting, poor generalization, and numerical instability (Hastie et al., 2009; Salehi et al., 2019).

Motivated by these limitations, this work proposes a scalable reformulation of logistic regression for feature selection in high-dimensional datasets, aiming to preserve interpretability while improving applicability in settings where the number of attributes substantially exceeds the number of samples.

## 5 Scalable Modified Logistic Regression for High-Dimensional Feature Selection

Let *A* ∈ ℝ^*N ×p*^ be the data matrix, where *N* denotes the number of samples and *p* denotes the number of attributes. Each row *x*_*j*_ ∈ ℝ^*p*^, *j* = 1, …, *N*, represents the attribute vector associated with the *j*-th sample. In the binary classification setting considered in this work, each sample is associated with a class label *y*_*j*_ ∈ {0, 1}, where the two values represent the control and disease groups, respectively.

In classical logistic regression, the probability that a sample *x*_*j*_ belongs to the positive class is modeled as

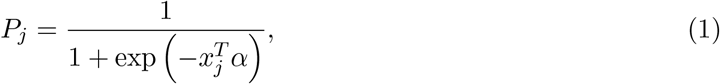

where *α* ∈ R^*p*^ is the vector of coefficients associated with the attributes. If an intercept term is required, it can be incorporated by adding a column of ones to the matrix *A*. The value *P*_*j*_ represents a probability and therefore satisfies *P*_*j*_ ∈ [0, 1].

The logit transformation associated with *P*_*j*_ is defined as

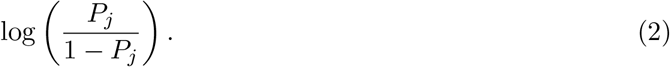

Applying this transformation to Equation 1, we obtain

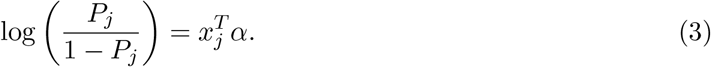

Thus, although the predicted probability is nonlinear, the logit of the probability is linear in the coefficient vector *α*. This property motivates the formulation adopted in this work.

To construct a linear representation of the binary classification problem, we encode the class labels through a log-odds transformation. Since the logit function is not defined for probabilities exactly equal to 0 or 1, we introduce a small constant *ε*, with 0 *< ε <* 0.5, and define a transformed response vector *b* ∈ ℝ^*N*^ as

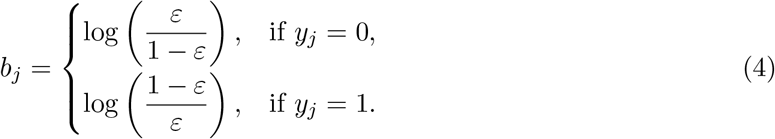

In all experiments, we set *ε* = 10^*−*5^. This value was chosen to avoid undefined logit values for class labels encoded as 0 or 1, while keeping the transformed labels close to the extreme class probabilities. Since the transformation maps the two classes to symmetric values,

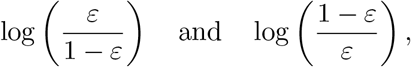

*ε* controls the scale of the transformed response vector *b*, but not the sign assigned to each class. In this formulation, samples from different classes are mapped to opposite regions of the log-odds scale. Therefore, the classification problem can be represented as the following linear inverse problem:

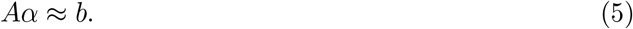

A direct way to estimate *α* would be to minimize the squared residual error,

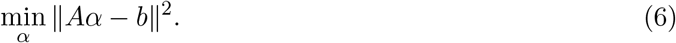

This is the classical least-squares formulation for an overdetermined or approximately consistent linear system. However, in high-dimensional biomedical datasets, the number of attributes is often much larger than the number of samples, that is, *p* ≫ *N* . This situation is common in gene expression analysis, where tens of thousands of molecular attributes may be measured for a relatively small number of patients.

In such scenarios, the least-squares problem may be underdetermined, leading to infinitely many possible solutions. Moreover, the matrix *A*^*T*^ *A* may be singular or ill-conditioned, making the estimation of *α* unstable and highly sensitive to small perturbations in the data. To obtain a stable solution, we introduce an objective function that minimizes both the residual error and the magnitude of the coefficient vector:

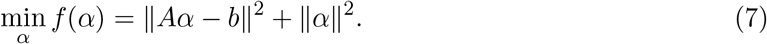

The first term in Equation 7 measures the discrepancy between the transformed response vector *b* and the linear predictor *Aα*. The second term penalizes large coefficient values, reducing instability and improving the conditioning of the estimation problem. This is especially important when the number of attributes substantially exceeds the number of samples.

The objective function can be written in matrix form as

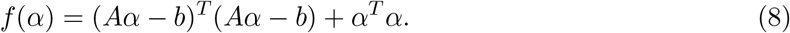

Expanding the first term, we obtain

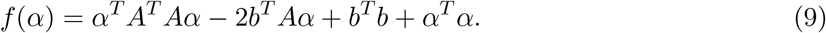

To find the minimizer of *f* (*α*), we compute the gradient with respect to *α*:

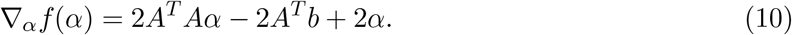

Setting the gradient equal to zero gives

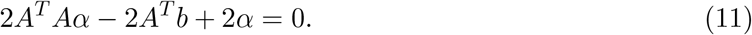

Dividing by 2, we obtain

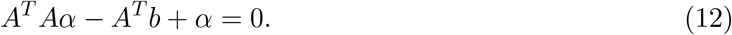

Therefore,

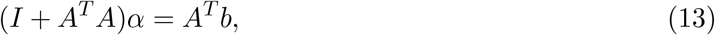

where *I* is the identity matrix of size *p* × *p*. Equation 13 characterizes the coefficient vector *α* that minimizes the objective function.

Although Equation 13 provides a direct mathematical characterization of the solution, explicitly forming *A*^*T*^ *A* or computing a matrix inverse may be computationally expensive and numerically undesirable in high-dimensional settings. To address this issue, we reformulate the problem as an equivalent augmented linear system:

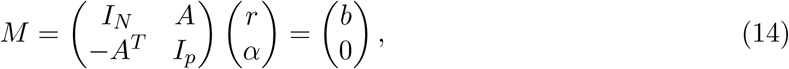

where *I*_*N*_ and *I*_*p*_ are identity matrices of sizes *N* × *N* and *p* × *p*, respectively, and *r* ∈ ℝ^*N*^ is an auxiliary variable.

The equivalence between Equation 14 and Equation 13 can be shown from the two block equations. From the first block row, we have

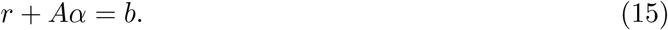

From the second block row, we have

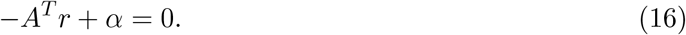

Thus,

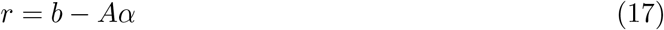

and

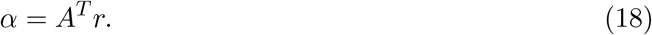

Substituting Equation 17 into Equation 18, we obtain

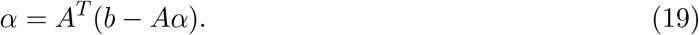

Therefore,

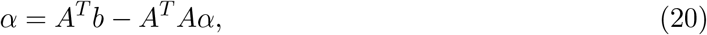

which leads to

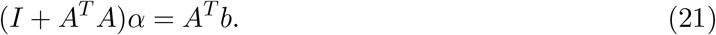

This is exactly the normal equation obtained from the minimization of Equation 7. Hence, the augmented system in Equation 14 is mathematically equivalent to the regularized formulation.

The advantage of the augmented formulation is computational. Instead of explicitly forming *A*^*T*^ *A* or computing a dense matrix inverse, the model can exploit the block structure and sparsity of the augmented matrix. This allows the use of sparse linear algebra techniques, such as sparse LU decomposition or iterative solvers, reducing memory usage and improving scalability for high-dimensional datasets.

After estimating *α*, the coefficients are used to rank the attributes according to their contribution to the separation between classes. Specifically, attributes with larger absolute coefficient values |*α*_*i*_| are interpreted as having greater influence on the classification boundary. Therefore, feature selection is performed by ranking the attributes according to |*α*_*i*_| and retaining the top *k* attributes with the largest absolute coefficients, |*α*_*i*_|. In the classification experiments, *k* = 20 was fixed for the proposed method in all datasets. This fixed value was chosen to provide a compact and comparable gene signature across datasets and to evaluate whether a small number of highly ranked attributes would be sufficient for discrimination between sepsis and control samples.

It is important to distinguish the number of features used in each classification model from the total number of unique genes reported in the stability analysis. In Table 1, the column “# Features” refers to the number of features retained within each cross-validation training fold for model fitting and test-fold prediction. Therefore, for the proposed method, each fold used the top 20 attributes ranked by |*α*_*i*_| from the corresponding training set. In contrast, Table 2 reports the union of unique genes selected across all cross-validation folds for each dataset. Because feature selection was repeated independently within each training fold, different folds could select partially different top-ranked genes. Consequently, the total number of unique selected genes across folds can be greater than 20, as observed.

**Table 1.**
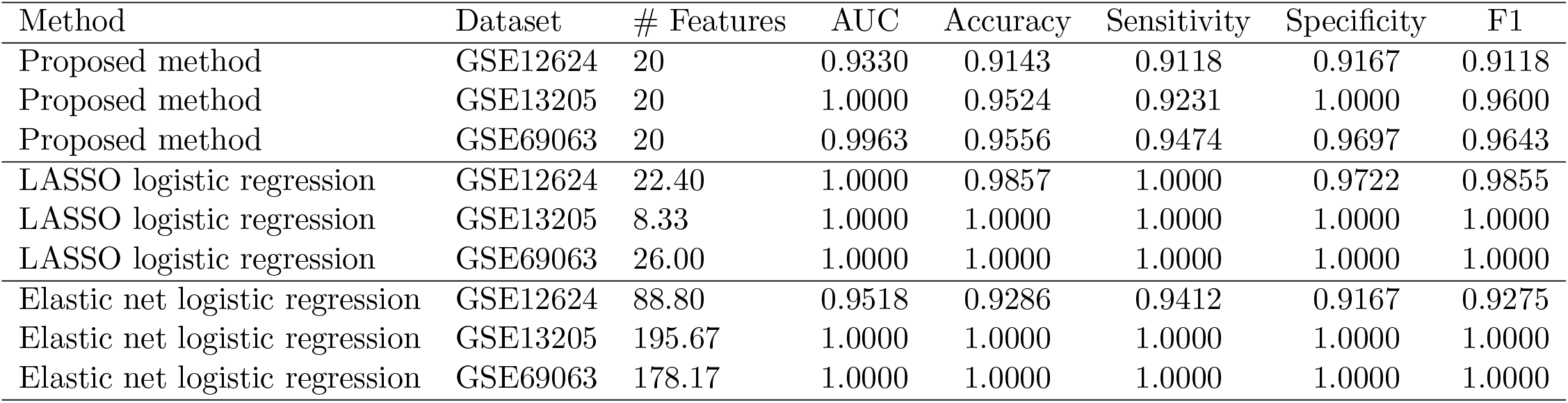
Classification performance comparison across the evaluated datasets. The column “# Features” indicates the number of features used in each cross-validation fold for model fitting and prediction. For the proposed method, the top 20 attributes ranked by |*α*_*i*_| were retained in each fold.

**Table 2.**
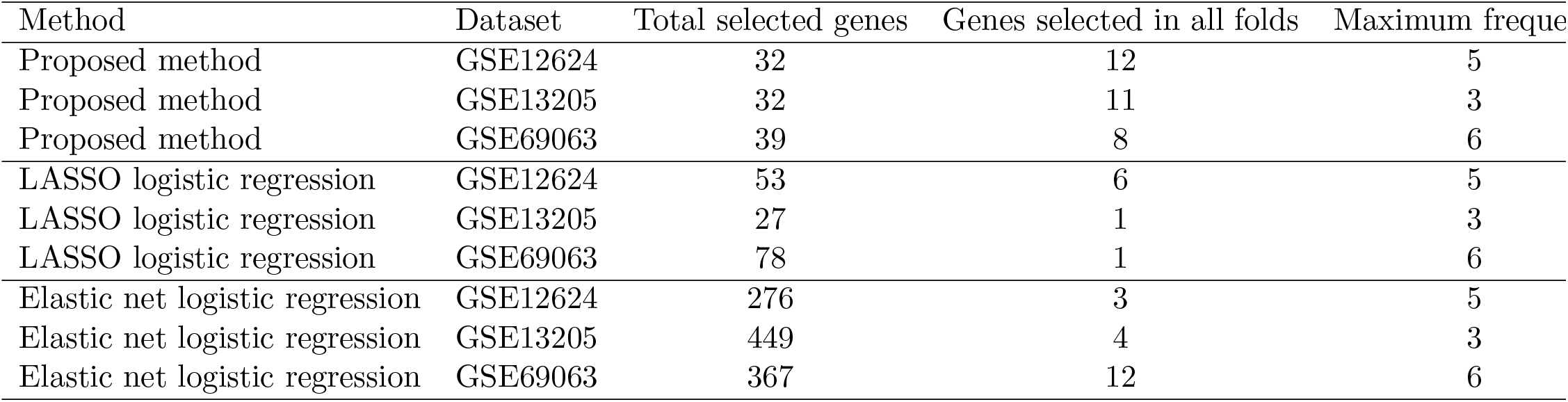
Stability of feature selection across cross-validation folds. “Total selected genes” corresponds to the number of unique genes selected at least once across all folds, whereas “Genes selected in all folds” indicates the subset of genes repeatedly selected in every fold of the corresponding dataset.

| Method | Dataset | Total selected genes | Genes selected in all folds | Maximum frequency |
| --- | --- | --- | --- | --- |
| Proposed method | GSE12624 | 32 | 12 | 5 |
| Proposed method | GSE13205 | 32 | 11 | 3 |
| Proposed method | GSE69063 | 39 | 8 | 6 |
| LASSO logistic regression | GSE12624 | 53 | 6 | 5 |
| LASSO logistic regression | GSE13205 | 27 | 1 | 3 |
| LASSO logistic regression | GSE69063 | 78 | 1 | 6 |
| Elastic net logistic regression | GSE12624 | 276 | 3 | 5 |
| Elastic net logistic regression | GSE13205 | 449 | 4 | 3 |
| Elastic net logistic regression | GSE69063 | 367 | 12 | 6 |

The proposed approach combines three elements: a log-odds encoding of the binary class labels, a linear formulation for coefficient estimation, and an equivalent sparse augmented system for scalable computation. This combination enables the application of a logistic-regression-inspired model to high-dimensional biomedical datasets, where the number of attributes is much larger than the number of samples, while preserving interpretability through the analysis of the estimated coefficients.

## 6 Results and Analysis

### 6.1 Experimental Evaluation Setup

All methods were evaluated using stratified cross-validation. The number of folds was defined according to the sample size and class distribution of each dataset, in order to preserve both classes in every training and test partition. Specifically, we used 5 folds for GSE12624, 3 folds for GSE13205, and 6 folds for GSE69063. The smaller number of folds in GSE13205 was adopted because this dataset contained only 21 samples, including 8 controls and 13 sepsis samples, making larger stratified partitions less stable. The use of dataset-specific fold numbers ensured that each test fold contained representative samples from both classes while reducing the risk of unstable estimates in the smallest dataset.

In each fold, the training set was used for all model-fitting steps, including feature ranking and feature selection. The corresponding test fold was kept completely independent from the feature selection process and was used only for performance evaluation. This design was adopted to avoid information leakage from the test samples into the feature selection step.

For the proposed method, the coefficient vector *α* was estimated using only the training data in each fold. Features were ranked by the absolute coefficient magnitude, |*α*_*i*_|, and the top 20 attributes were retained for classification. The same training/test splits were used for the proposed method, LASSO logistic regression, and elastic net logistic regression to ensure comparability across methods.

Class imbalance was handled through stratified partitioning, preserving the approximate proportion of sepsis and control samples in each fold. No additional resampling procedure, such as oversampling or undersampling, was applied. A fixed random seed of 42 was used to make the fold assignments reproducible.

For the baseline methods, LASSO and elastic net logistic regression were fitted using only the training data within each cross-validation fold. Regularization parameters were selected using the default internal cross-validation procedure of the implementation used in this study. No information from the test fold was used during model fitting, feature selection, or hyperparameter selection. Nevertheless, because an explicitly nested validation scheme with independently defined tuning grids was not performed, comparative performance results should be interpreted with caution.

### 6.2 Coefficient Distribution and Feature Selection

The coefficient vector *α* estimated by the proposed method was used to rank the genes according to their contribution to class separation. Figures 1, 2, and 3 show the ordered *α* coefficients obtained for the three sepsis datasets.

**Figure 1.**
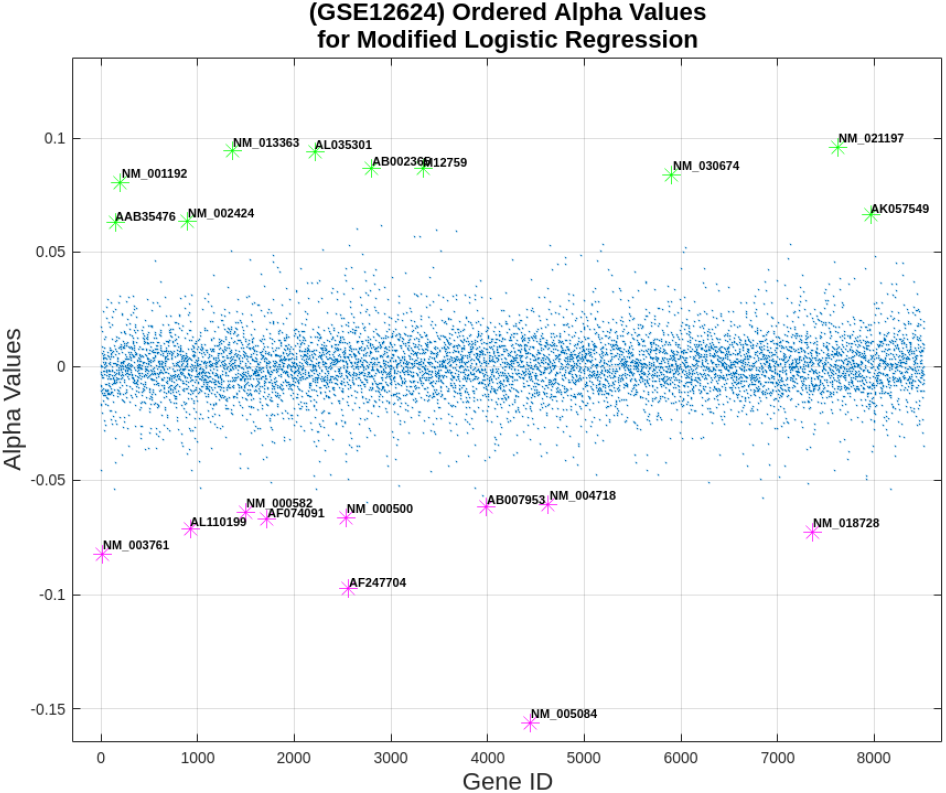
Ordered *α* coefficients estimated by the proposed method. Highlighted points indicate the attributes with the largest absolute coefficients, |*α*_*i*_|, used for feature selection.

**Figure 2.**
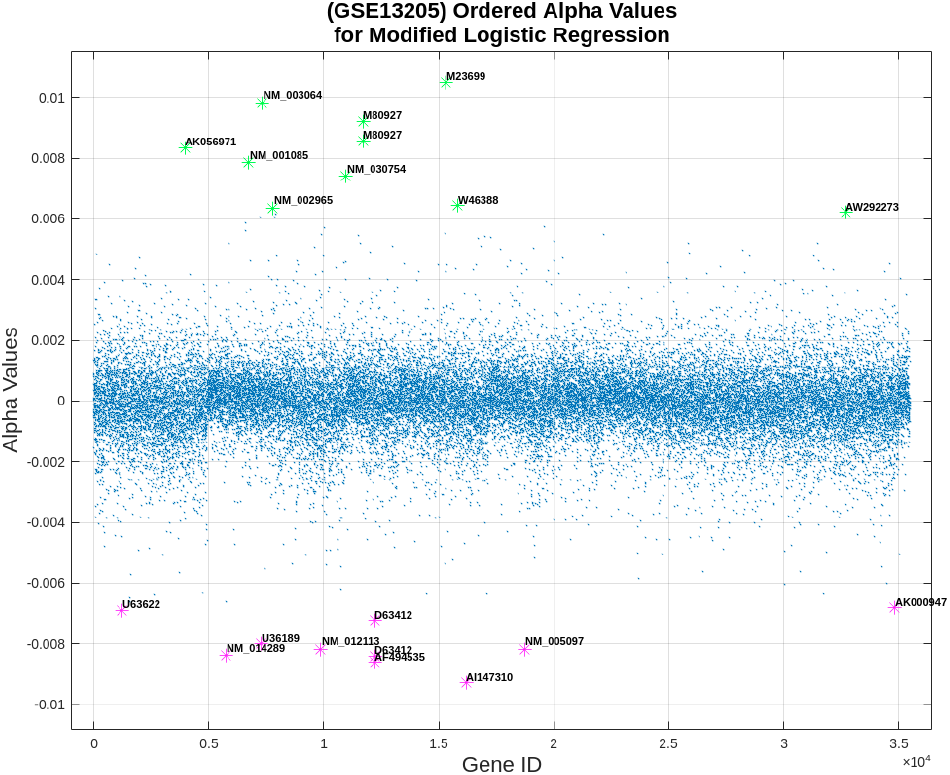
Ordered *α* coefficients estimated by the proposed method. Highlighted points indicate the attributes with the largest absolute coefficients, |*α*_*i*_|, used for feature selection.

**Figure 3.**
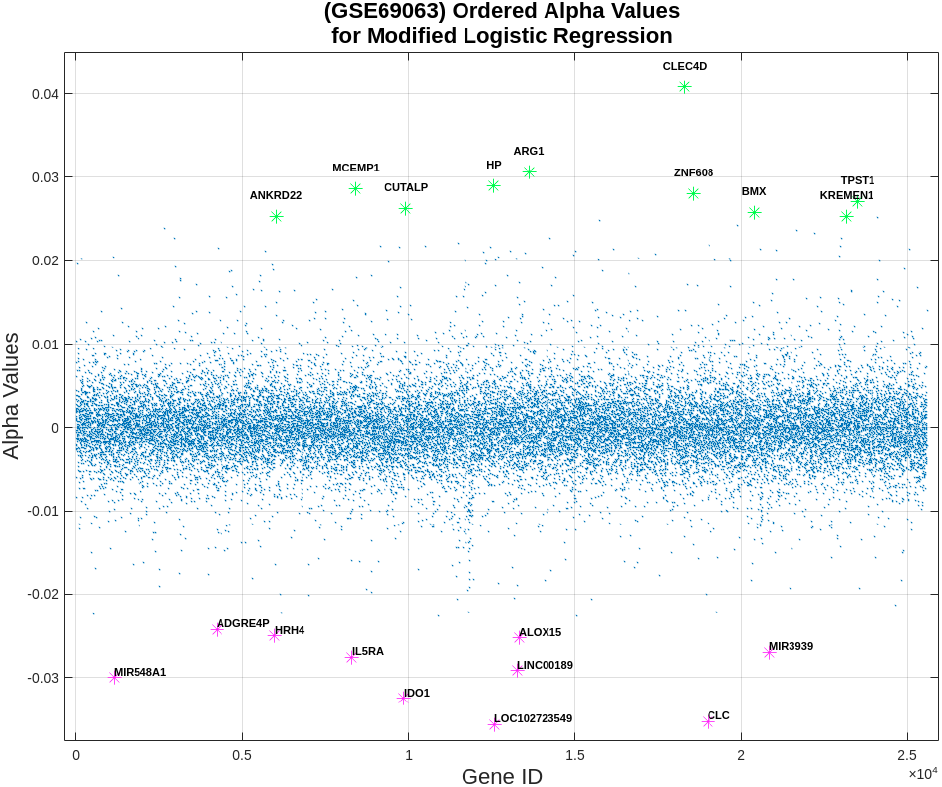
Ordered *α* coefficients estimated by the proposed method. Highlighted points indicate the attributes with the largest absolute coefficients, |*α*_*i*_|, used for feature selection.

Across the three datasets, most coefficients were concentrated around zero, indicating that the majority of genes had limited influence on the separation between control and sepsis samples. In contrast, a small subset of genes presented larger positive or negative coefficient values. These genes were considered the most relevant attributes and were selected for subsequent classification and analysis.

This coefficient distribution supports the use of the proposed method as a feature selection strategy. By ranking genes according to |*α*_*i*_|, the method identifies a reduced subset of attributes with stronger discriminatory relevance, while preserving the interpretability of the model through the analysis of the estimated coefficients.

### 6.3 SVD Visualization Before and After Feature Selection

To further examine the effect of feature selection on the structure of the data, we used singular value decomposition (SVD) to visualize the samples before and after selecting the genes with the largest |*α*_*i*_| values. SVD is widely used for dimensionality reduction and exploratory visualization of high-dimensional data, and has also been applied to genome-wide expression data analysis (Alter et al., 2000). Figure 4 shows the projections obtained for the three sepsis datasets.

**Figure 4.**
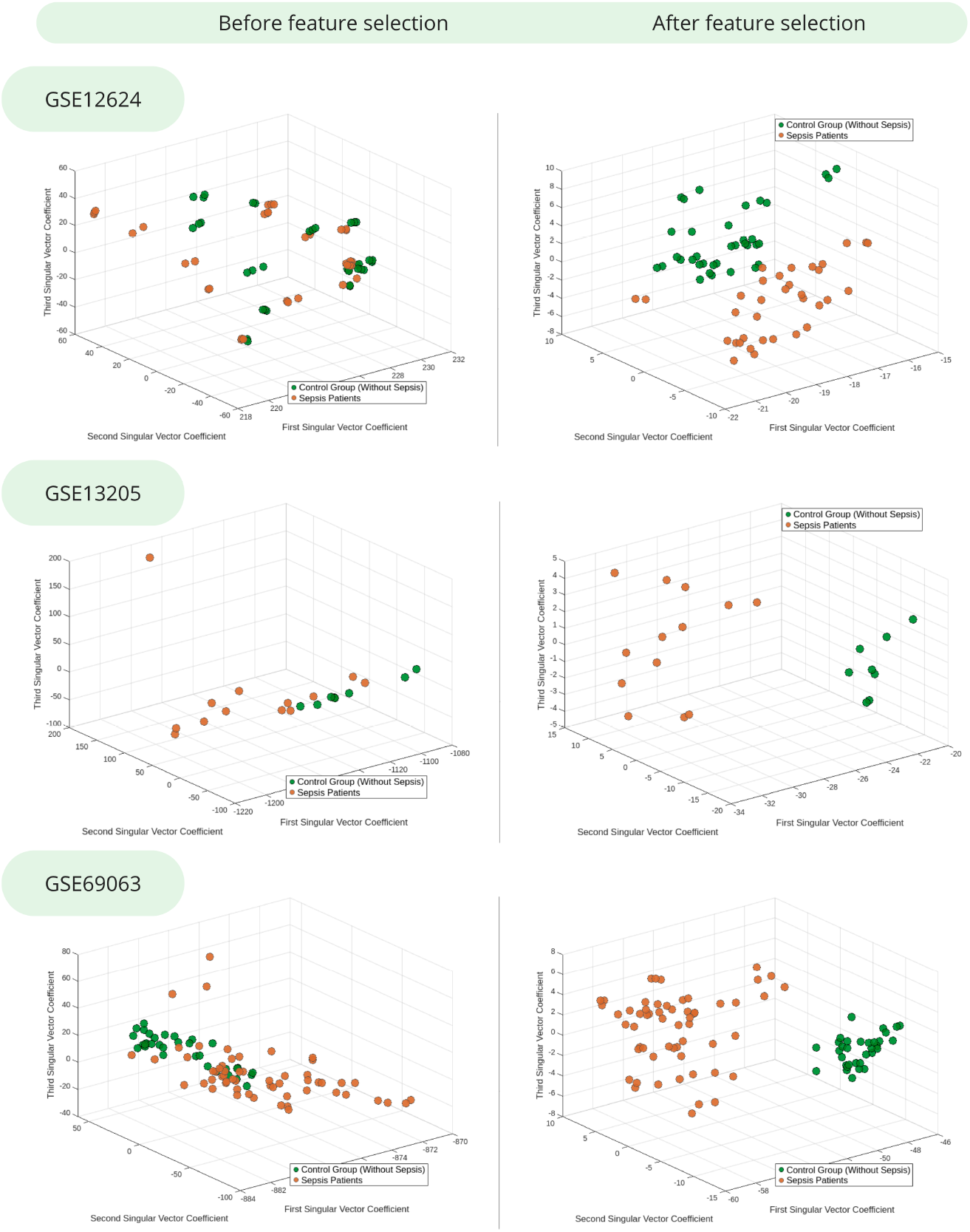
SVD visualization before and after feature selection. After selecting attributes with the largest |*α*_*i*_|, control and sepsis samples show a clearer separation pattern.

Before feature selection, control and sepsis samples showed a higher degree of overlap in the projected space, indicating that the original high-dimensional representation contained substantial variability not directly related to class separation. After retaining only the genes selected by the proposed method, a clearer separation pattern between the two groups was observed in Figure 4.

This result suggests that the selected genes preserve relevant discriminatory information while reducing the dimensionality of the original data. However, the SVD projection should be interpreted as an exploratory visualization rather than as a formal measure of classification performance.

Quantitative metrics obtained through cross-validation are therefore used in the following sections to assess predictive performance more rigorously.

### 6.4 Classification Performance

Table 1 summarizes the classification performance obtained by the evaluated methods across the three sepsis datasets. Figures 5 and 6 provide a visual comparison of the AUC and F1-score values, respectively.

**Figure 5.**
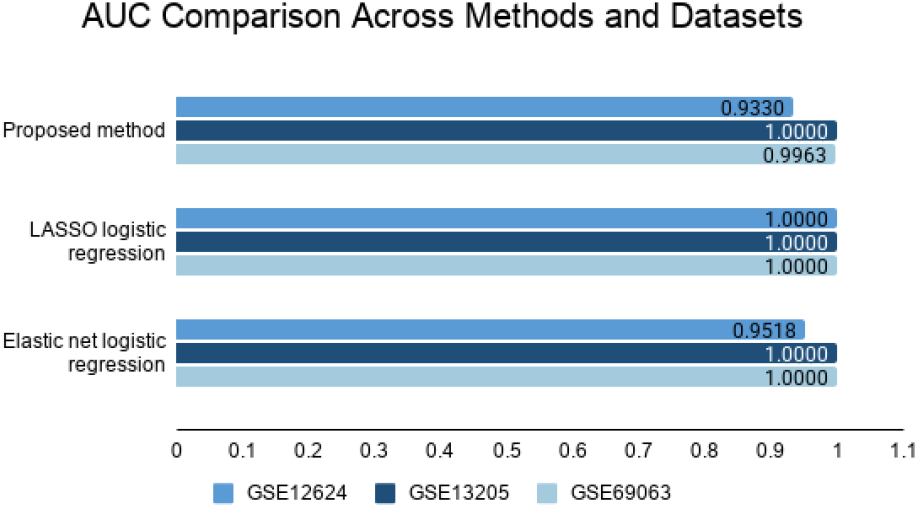
AUC comparison across the evaluated methods and datasets.

**Figure 6.**
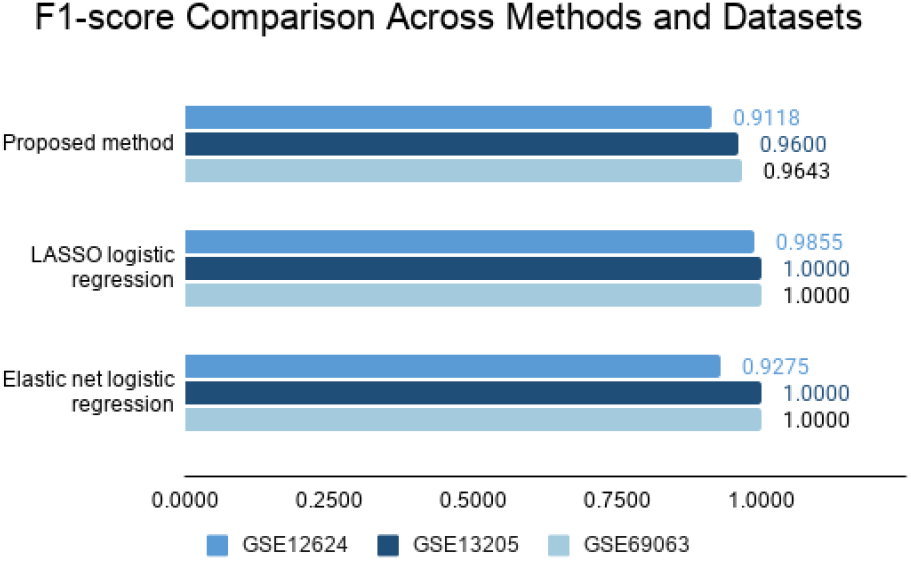
F1-score comparison across the evaluated methods and datasets.

The proposed method achieved high predictive performance in all datasets. In GSE12624, it obtained an AUC of 0.9330 and an F1-score of 0.9118. In GSE13205, the method achieved an AUC of 1.0000 and an F1-score of 0.9600. In GSE69063, it achieved an AUC of 0.9963 and an F1-score of 0.9643. These results indicate that the selected genes retained relevant information for distinguishing control and sepsis samples.

LASSO logistic regression achieved the highest overall predictive performance, with an AUC of 1.0000 in the three datasets. It also obtained perfect F1-score values in GSE13205 and GSE69063. Elastic net logistic regression also showed strong performance, with an AUC of 0.9518 in GSE12624 and an AUC of 1.0000 in both GSE13205 and GSE69063. These results confirm that logistic regression models are strong baselines for high-dimensional gene expression classification.

Overall, the proposed method did not always outperform LASSO or elastic net in terms of predictive performance alone. Nevertheless, it achieved competitive classification results while producing compact and interpretable gene signatures. Therefore, the predictive results should be interpreted together with the feature selection stability and parsimony analyses presented in the following sections.

### 6.5 Feature Selection Stability

Table 2 summarizes the stability and parsimony of the gene sets selected by each method across the cross-validation folds. The reported values correspond to the averages obtained across the cross-validation runs, reflecting the mean behavior of each method rather than the result of a single execution. Figure 7 provides a visual comparison of the average total number of selected genes for each method and dataset.

**Figure 7.**
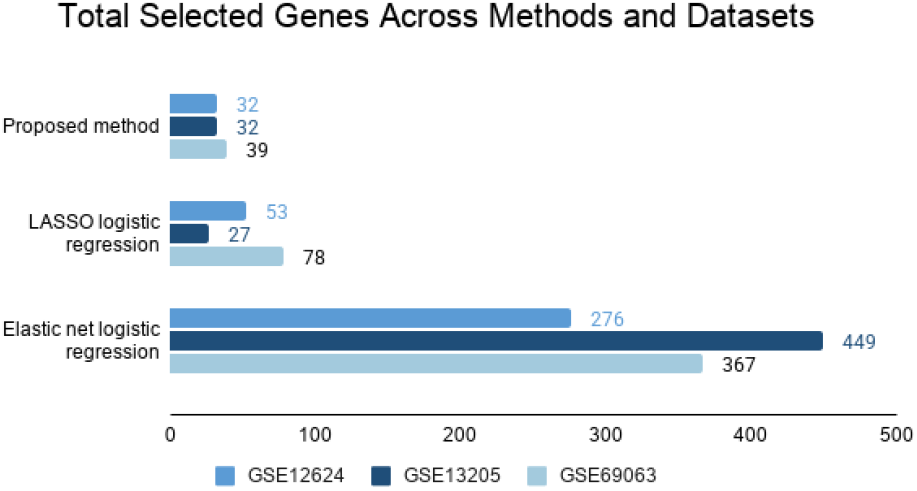
Total number of selected genes across cross-validation folds for each method and dataset.

The proposed method selected compact gene signatures in the three datasets, with 32 genes in GSE12624, 32 genes in GSE13205, and 39 genes in GSE69063. In comparison, LASSO selected 53 genes in GSE12624, 27 genes in GSE13205, and 78 genes in GSE69063, while elastic net selected substantially larger sets, with 276 genes in GSE12624, 449 genes in GSE13205, and 367 genes in GSE69063. These results indicate that the proposed method tends to produce more compact signatures than elastic net and, in most datasets, also fewer selected genes than LASSO.

The stability of the selected genes was also evaluated by counting how many genes were selected in all cross-validation folds. The proposed method selected 12 genes in all folds for GSE12624, 11 genes in all folds for GSE13205, and 8 genes in all folds for GSE69063. This suggests that part of the selected signature remained stable despite changes in the training set. However, the number of folds differs across datasets, with maximum frequencies of 5, 3, and 6 for GSE12624, GSE13205, and GSE69063, respectively. Therefore, stability should be interpreted within each dataset rather than by directly comparing raw frequencies across datasets.

Although LASSO and elastic net achieved higher predictive performance in some cases, they often selected larger gene sets, especially elastic net. This reflects a trade-off between predictive performance and parsimony. In this context, the proposed method provides a compact and stable alternative for feature selection, which may be advantageous when biological interpretation and the identification of a reduced set of candidate genes are important objectives.

### 6.6 Computational Performance

Table 3 summarizes the computational performance of the evaluated methods, and Figure 8 shows the total runtime of the cross-validation procedure for each method and dataset.

**Table 3.** Computational performance of the evaluated methods.

| Method | Dataset | Training time (s) | Prediction time (s) | Total time (s) |
| --- | --- | --- | --- | --- |
| Proposed method | GSE12624 | 0.565161 | 0.001054 | 0.620734 |
| Proposed method | GSE13205 | 1.622316 | 0.000036 | 1.637542 |
| Proposed method | GSE69063 | 4.646431 | 0.00008 | 4.690719 |
| LASSO logistic regression | GSE12624 | 53.098582 | 0.001299 | 53.117219 |
| LASSO logistic regression | GSE13205 | 6.463061 | 0.000677 | 6.478521 |
| LASSO logistic regression | GSE69063 | 8.311969 | 0.001849 | 8.360391 |
| Elastic net logistic regression | GSE12624 | 124.458177 | 0.001493 | 124.476351 |
| Elastic net logistic regression | GSE13205 | 4.793038 | 0.000425 | 4.806646 |
| Elastic net logistic regression | GSE69063 | 5.076891 | 0.001554 | 5.124437 |

**Figure 8.**
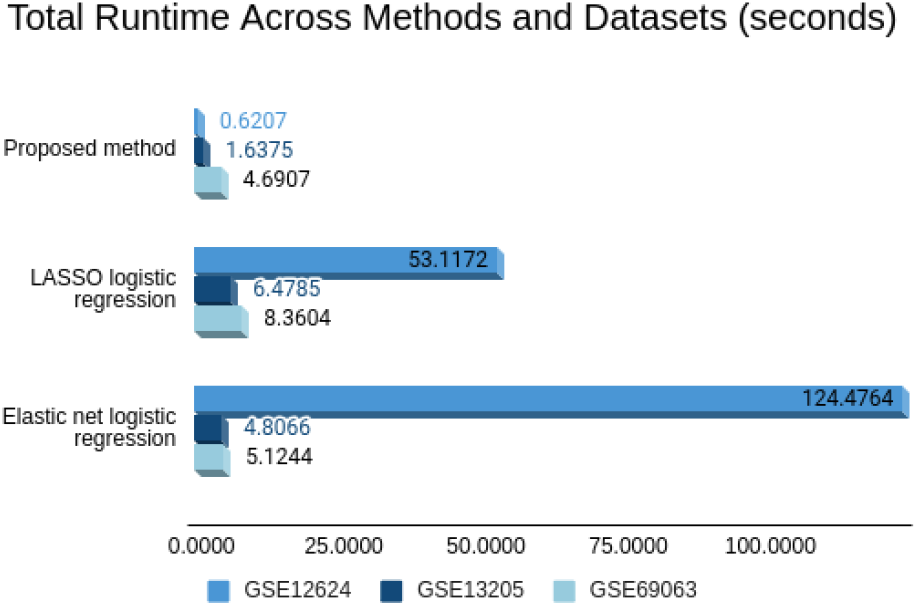
Total runtime of the cross-validation procedure for each evaluated method and dataset in seconds.

The proposed method presented low to moderate computational cost across the three datasets. The total runtime was 0.6207 seconds for GSE12624, 1.6375 seconds for GSE13205, and 4.6907 seconds for GSE69063. These values indicate that the proposed formulation can be applied efficiently to high-dimensional gene expression datasets, even when the number of attributes is substantially larger than the number of samples.

Compared with the regularized logistic regression baselines, the proposed method showed substantially lower runtime in GSE12624. In this dataset, LASSO logistic regression required 53.1172 seconds, while elastic net logistic regression required 124.4764 seconds. This difference suggests that the proposed linear-system-based formulation may reduce computational cost in settings where penalized logistic regression requires longer optimization times.

For GSE13205 and GSE69063, the differences in runtime were less pronounced, and the relative computational advantage varied across methods. LASSO and elastic net remained computationally feasible in these datasets, although their selected gene sets were generally larger than those produced by the proposed method. Therefore, computational performance should be interpreted together with the number of selected genes and the feature selection behavior of each method.

Overall, these results indicate that the proposed method provides competitive computational performance, especially when compared with regularized logistic regression models, while maintaining a compact and interpretable feature selection strategy.

### 6.7 Biological Interpretation of Selected Genes

Although an in-depth biological validation of the selected genes is beyond the main scope of this study, we performed an exploratory biological interpretation of a subset of attributes selected by the proposed method. This subset was defined from the genes highlighted by the coefficient-based ranking, that is, genes with larger absolute values of |*α*_*i*_| in each dataset, together with candidates that appeared biologically plausible or had previous support in sepsis-related transcriptomic or biomarker studies.

Therefore, Table 4 does not represent an exhaustive list of all selected genes across the cross-validation folds. Instead, it summarizes representative candidates selected by the proposed method that were considered relevant for biological interpretation. For each dataset, we included genes that either had prior evidence in the sepsis literature or were associated with biological processes compatible with sepsis pathophysiology, such as immune and inflammatory responses, leukocyte activation, cytokine and receptor-mediated signaling, metabolic regulation, oxidative stress, and tissue remodeling.

**Table 4.** Exploratory biological relevance of representative attributes selected by the proposed method.

| Dataset | Attribute | Status | Reason for inclusion | Supporting evidence and biological role |
| --- | --- | --- | --- | --- |
| GSE12624 | MMP8 | Reported | Highlighted by the coefficient-based ranking and previously reported in sepsis studies. | Previously reported in sepsis studies (Zhang et al., 2023; Lyu et al., 2025; Kong et al., 2023; Xiao and Zhang, 2024; Zhang et al., 2024); related to neutrophil-associated inflammation and extracellular matrix remodeling. |
| GSE12624 | CXCL10 | Reported | Highlighted by the coefficient-based ranking and previously reported in sepsis studies. | Previously reported in sepsis studies (Xie, 2022; He et al., 2025); involved in chemokine-mediated immune and inflammatory signaling. |
| GSE12624 | PCOLCE2 | Exploratory | Selected by the model and retained as a biologically plausible candidate related to tissue remodeling. | No direct supporting sepsis reference was assigned in this exploratory table; biologically interpreted in relation to extracellular matrix organization and collagen-associated tissue remodeling. |
| GSE12624 | WFDC1 | Exploratory | Selected by the model and retained as a biologically plausible candidate related to inflammatory regulation. | No direct supporting sepsis reference was assigned in this exploratory table; biologically interpreted in relation to tissue remodeling and inflammatory regulation. |
| GSE13205 | IL6R | Exploratory | Highlighted by the coefficient-based ranking and functionally related to cytokine receptor signaling. | No direct supporting sepsis reference was assigned in this exploratory table; biologically interpreted in relation to cytokine receptor signaling and immune/inflammatory response. |
| GSE13205 | SLPI | Reported | Highlighted by the coefficient-based ranking and previously reported in sepsis studies. | Previously reported in sepsis studies (Tang et al., 2021; Lu et al., 2022; Chen et al., 2025); related to antimicrobial response, inflammation regulation, and tissue protection. |
| GSE13205 | CHI3L1 | Reported | Highlighted by the coefficient-based ranking and previously reported in sepsis studies. | Previously reported in sepsis studies (Van Der Aart et al., 2025); related to inflammatory response and extracellular matrix/tissue remodeling. |
| GSE13205 | AQP4 | Exploratory | Selected by the model and retained due to tissue-specific transcriptomic discrimination. | No direct supporting sepsis reference was assigned in this exploratory table; biologically interpreted in relation to water transport and tissue-specific transcriptomic discrimination. |
| GSE69063 | ARG1 | Reported | Highlighted by the coefficient-based ranking and previously reported in sepsis studies. | Previously reported in sepsis studies (Zhang et al., 2022; Dai et al., 2022; Winkler et al., 2017); related to arginine metabolism, immune regulation, and metabolic reprogramming. |
| GSE69063 | HP | Reported | Highlighted by the coefficient-based ranking and previously reported in sepsis studies. | Previously reported in sepsis studies (Lyu et al., 2025; Kong et al., 2023; Van Der Aart et al., 2025); related to acute-phase response, inflammation, and oxidative-stress-associated processes. |
| GSE69063 | MCEMP1 | Reported | Highlighted by the coefficient-based ranking and previously reported in sepsis studies. | Previously reported in sepsis studies (Lyu et al., 2025; Kong et al., 2023); related to myeloid-cell activation and innate immune response. |
| GSE69063 | IDO1 | Reported | Highlighted by the coefficient-based ranking and previously reported in sepsis studies. | Previously reported in sepsis studies (Changsirivathanathamrong et al., 2011; Darcy et al., 2011; Stier et al., 2025); related to tryptophan metabolism and immunoregulatory response. |
| GSE69063 | ALOX15 | Exploratory | Selected by the model and retained as a biologically plausible candidate related to lipid and redox pathways. | No direct supporting sepsis reference was assigned in this exploratory table; biologically interpreted in relation to lipid metabolism, oxidative stress, and inflammatory regulation. |

Among the genes highlighted by the model, candidates such as *IL6R, CXCL10, ARG1, IDO1, CHI3L1, MMP8, HP, SLPI*, and *MCEMP1* are particularly noteworthy because they are related to processes commonly implicated in sepsis, including immune regulation, inflammatory signaling, neutrophil activity, extracellular matrix remodeling, and metabolic or oxidative-stress-related pathways. Some of these genes have already been reported in sepsis transcriptomic or biomarker studies, whereas others appear to represent less explored or context-dependent signals.

Thus, the selected gene sets should not be interpreted as definitive biomarkers, but rather as exploratory candidates with potential biological relevance. The main objective of this work remains the evaluation of a modified logistic-regression-based strategy for feature selection in high-dimensional gene expression data. Nevertheless, the biological plausibility of several selected genes reinforces the interpretability of the proposed method and supports its potential use as a computational tool for identifying compact and meaningful gene signatures in transcriptomic studies.

The *Status* column in Table 4 provides an exploratory indication of whether each selected gene has already been reported in previous sepsis-related transcriptomic or biomarker studies. Genes labeled as *Reported* correspond to candidates with prior evidence in the sepsis literature, whereas genes labeled as *Exploratory* refer to candidates selected by the proposed method that showed biological plausibility but appear to be less frequently described or more context-dependent in previous sepsis biomarker studies. The column *Reason for inclusion* clarifies why each gene was retained for biological discussion, linking the table back to the coefficient-based feature selection step. Therefore, the table should not be interpreted as definitive biomarker validation, but as a qualitative summary of representative selected genes with potential biological relevance.

### 6.8 Overall Discussion and Limitations

The comparative analysis shows that LASSO and elastic net logistic regression achieved the highest predictive performance in some of the evaluated datasets. In particular, these models obtained perfect or near-perfect classification results in several cases. However, this performance was often associated with larger selected gene sets, especially for elastic net. In contrast, the proposed method achieved competitive classification performance while producing more compact gene signatures, which may be advantageous when interpretability and biological analysis are central objectives.

The SVD visualizations provide additional exploratory evidence that the genes selected by the proposed method retain relevant discriminatory information. After feature selection, control and sepsis samples showed a clearer separation pattern in the projected space. Although this visualization does not replace quantitative validation, it supports the interpretation that the selected genes capture class-related structure in the data.

The stability analysis further supports the relevance of the selected signatures. Several genes were repeatedly selected across all cross-validation folds, suggesting that the feature ranking obtained by the proposed method was not entirely dependent on a specific training partition. This recurrence is important in high-dimensional biomedical data, where unstable feature selection may lead to gene lists that are difficult to interpret or reproduce.

Despite these findings, some limitations should be acknowledged. First, the datasets used in this study have relatively small sample sizes, which is common in gene expression studies but increases the risk of overfitting and limits the generalizability of the results. Second, the datasets may differ in terms of platform, cohort composition, preprocessing, and biological context, which can affect both classification performance and gene selection. Third, no independent external validation cohort was used, and therefore the generalization of the selected gene signatures to new datasets remains to be further investigated.

In addition, the biological interpretation of the selected genes should be considered preliminary. Although some selected genes have been previously associated with sepsis or immune-related mechanisms, the remaining candidates require further validation through independent datasets and experimental studies. Finally, the perfect performance observed for some baseline methods may reflect strong separability in specific datasets, but it may also indicate overfitting or sensitivity to small sample sizes. For this reason, predictive performance should be interpreted together with feature stability, parsimony, and biological plausibility.

Some methodological choices should also be considered when interpreting the results. First, the proposed method requires the specification of the logit-scaling parameter *ε*. In this study, we used a fixed value of *ε* = 10^*−*5^ to avoid undefined logit transformations for labels encoded as 0 and 1. However, we did not perform a formal sensitivity analysis across alternative values of *ε*. Future work should evaluate whether different choices of *ε* affect coefficient magnitudes, gene ranking stability, and selected signatures.

Second, the number of selected features was fixed at 20 for the classification experiments. This choice provided compact and comparable signatures across datasets, but other values of *k* may affect both predictive performance and biological interpretation. Third, the total number of selected genes reported in the stability analysis corresponds to the union of genes selected across folds, not to the number of genes used in each individual model. Therefore, these values should be interpreted as indicators of feature selection variability across cross-validation partitions.

Overall, the proposed method should not be interpreted as consistently outperforming regularized logistic regression in predictive accuracy. Rather, it provides a competitive and interpretable alternative for feature selection in high-dimensional gene expression data, offering a balance between classification performance, compact gene signatures, computational cost, and biological interpretability.

## 7 Conclusion

In this work, we proposed a scalable modified logistic regression framework for feature selection in high-dimensional gene expression data. The method combines a log-odds encoding of binary class labels with a linear formulation and an equivalent augmented linear system, allowing coefficient estimation without explicitly computing dense matrix inverses. By ranking genes according to the absolute magnitude of the estimated coefficients, the proposed approach provides a simple and interpretable mechanism for selecting relevant attributes in settings where the number of features greatly exceeds the number of samples.

The method was evaluated on three publicly available sepsis gene expression datasets: GSE12624, GSE13205, and GSE69063. The results showed that the proposed approach achieved high classification performance across all datasets while selecting compact gene signatures. Although LASSO and elastic net logistic regression achieved higher or perfect predictive performance in some cases, they often selected larger sets of genes, particularly elastic net. Therefore, the proposed method should not be interpreted as a replacement for regularized logistic regression in terms of predictive accuracy alone, but rather as a competitive alternative when interpretability, parsimony, and computational simplicity are important objectives.

The analysis of feature selection stability showed that several genes were repeatedly selected across cross-validation folds, suggesting that the proposed coefficient-based ranking captured relevant discriminatory information. In addition, the SVD visualizations indicated that the selected genes improved the separation between control and sepsis samples in a reduced-dimensional representation.

Despite these promising results, some limitations should be considered. The evaluated datasets have relatively small sample sizes, which may limit the generalizability of the findings and increase the risk of overfitting. In addition, no independent external validation cohort was used, and the biological interpretation of the selected genes remains preliminary. Future work should evaluate the proposed method on larger and independent cohorts, investigate its robustness across different preprocessing strategies and omics platforms, and further explore the biological relevance of the selected genes.

Overall, the proposed framework provides an interpretable and computationally feasible approach for feature selection in high-dimensional gene expression data. Its ability to produce compact gene signatures while maintaining competitive classification performance suggests that it may be useful in biomedical applications.

## Data Availability

The datasets analyzed in this study are publicly available in the NCBI Gene Expression Omnibus under accession numbers GSE12624, GSE13205, and GSE69063. No new primary biological data were generated in this study.

## Code Availability

The source code used to implement the proposed method, perform the cross-validation experiments, generate the results, and reproduce the figures is publicly available at: https://github.com/rossanaoliveirasouza/modified_logistic_regression_method

The source code is provided as MATLAB scripts and functions in .m format and should be executed using MATLAB.

## Author Contributions

Rossana O. Souza: Conceptualization, Methodology, Software, Formal Analysis, Investigation, Data Curation, Visualization, Writing – Original Draft, and Writing – Review and Editing.

Wellington Francisco Rodrigues: Biological Interpretation, Validation, and Writing – Review and Editing.

Bráulio R. G. M. Couto: Validation, Supervision, and Writing – Review and Editing.

Marcos A. dos Santos: Conceptualization, Methodology, Supervision, Project Administration, and Writing – Review and Editing.

All authors read and approved the final version of the manuscript.

## Ethics Statement

This study was based exclusively on publicly available, de-identified datasets obtained from the NCBI Gene Expression Omnibus. No new participants were recruited, no identifiable personal data were accessed, and no new biological samples were collected. Therefore, additional institutional ethics approval and informed consent were not required for the present secondary analysis.

